# DLC1 is a direct target of activated YAP/TAZ that drives collective migration and sprouting angiogenesis

**DOI:** 10.1101/755389

**Authors:** Miesje van der Stoel, Lilian Schimmel, Kalim Nawaz, Anne-Marieke van Stalborch, Annett de Haan, Alexandra Klaus-Bergmann, Erik T. Valent, Duco S. Koenis, Geerten P. van Nieuw Amerongen, Carlie J. de Vries, Vivian de Waard, Martijn Gloerich, Jaap D. van Buul, Stephan Huveneers

## Abstract

YAP/TAZ signaling is crucial for sprouting angiogenesis and vascular homeostasis through the regulation of endothelial remodeling. Thus far the underlying molecular mechanisms that explain how YAP/TAZ control the vasculature remain unclear. We here identify Deleted-in-Liver-Cancer-1 (DLC1) as a direct transcriptional target of the activated YAP/TAZ-TEAD complex in the endothelium. Substrate stiffening and VEGF stimuli promote the endothelial expression of DLC1. DLC1 expression is dependent on the presence of YAP and TAZ, and constitutive activation of YAP efficiently promotes expression of DLC1. We show that DLC1 limits F-actin fiber formation, integrin-based focal adhesion lifetime and integrin-mediated traction forces. Depletion of endothelial DLC1 strongly perturbs cell polarization in directed collective migration and inhibits the formation of angiogenic sprouts. Importantly, the inability of YAP-depleted endothelial cells to collectively migrate and form angiogenic sprouts can be rescued by ectopic expression of DLC1. Together, these findings reveal that DLC1 fills a hitherto missing link between YAP/TAZ signaling and endothelial dynamics during angiogenesis.

## Introduction

The formation of new blood and lymphatic vessels through angiogenesis is essential for development and vital for tissue regeneration and tumorigenesis (Gomez-Salinero and Rafii, 2018; Potente et al., 2011). The luminal side of the vasculature is covered by a well-organized layer of endothelial cells. Angiogenesis is driven by endothelial cell proliferation and migration, during which the endothelial cells coordinate their movements collectively through tight remodeling of interactions with the vascular microenvironment and contacts between the endothelial cells (Betz et al., 2016; Szymborska and Gerhardt, 2018).

Yes-associated protein (YAP) and transcriptional co-activator with PDZ-binding motif (TAZ or WWTR1) proteins are key molecular switches shuttling between the cytoplasm and nucleus to control proliferation and migration (Panciera et al., 2017). Several studies have demonstrated the importance of YAP/TAZ for angiogenesis and vascular homeostasis (Choi et al., 2015; Nakajima et al., 2017a; Wang et al., 2016a, 2017; Neto et al., 2018). Mechanical cues such as extracellular matrix (ECM) stiffness and shear stress control the activity of YAP and TAZ (Dupont, 2016) and are important tissue properties that guide angiogenesis (Dorland and Huveneers, 2017; Choi et al., 2015; Nakajima et al., 2017b; Wang et al., 2016a, 2017; Neto et al., 2018). In addition, angiogenic signaling through vascular endothelial growth factor (VEGF) promotes activation of YAP/TAZ and a migratory transcriptional program to support developmental angiogenesis (Wang et al., 2017). YAP and TAZ activation is further regulated by Rho GTPase signaling and cytoskeletal contractility (Dupont et al., 2011; Elosegui-Artola et al., 2017).

Inactive YAP and TAZ are localized in the cytoplasm, whereas active YAP and TAZ (i.e. upon ECM stiffening, disturbed flow, or sparse cell densities) translocate to the nucleus (Dupont et al., 2011). Nuclear YAP/TAZ act as co-activators of TEA domain family members (TEAD) transcription factors to promote vascular development (Vassilev et al., 2001; Astone et al., 2018). Recently, it was shown that YAP/TAZ activation is needed to provide transcriptional feedback for collective migration of endothelial cells (Mason et al., 2019). Strikingly, the transcriptional target(s) of YAP/TAZ that are responsible for endothelial migration in angiogenesis remain to be identified.

We previously observed that Deleted-in-Liver-Cancer-1 (DLC1, also known as STARD12 or ARHGAP7) expression is high in endothelial cells on stiff substrates (Schimmel et al., 2018), pointing towards a downstream role for DLC1 in YAP/TAZ signaling. DLC1 is crucial for embryonic development and its depletion in mice leads to severe defects of various organs at embryonic day 10.5 (Durkin et al., 2005). DLC1 is an endothelial-enriched GTPase activating protein (GAP) that inactivates Rho-GTPases (van Buul et al., 2014). In addition, DLC1 has a serine-rich region that contains binding motifs for components of integrin-based focal adhesions (Barras and Widmann, 2014; Kim et al., 2009). Focal adhesions are crucial structures for mechanotransduction and migration as they connect cells to the ECM and mechanically couple the contractile actin cytoskeleton to the extracellular microenvironment (Geiger et al., 2001; Gardel et al., 2010; Grashoff et al., 2010).

In this study we identify DLC1 as a novel transcriptional target of the activated YAP/TAZ-TEAD complex. Expression of DLC1 is needed for integrin-based focal adhesion disassembly, cell polarization, collective cell migration and angiogenic sprouting. We further demonstrate that ectopic expression of DLC1 in YAP-depleted endothelial cells restores their migration and angiogenic sprouting capacity. In conclusion, we demonstrate that DLC1 is a crucial transcriptional target of YAP/TAZ in the endothelium. These findings place DLC1 as a key player in YAP/TAZ signaling and likely has wider implications for YAP/TAZ-driven flow sensing and the development of stiffness-related vascular diseases.

## Results

### DLC1 is a transcriptional target of YAP/TAZ and TEAD

We recently showed that DLC1 protein expression is high within stiff microenvironments of the vasculature (Schimmel et al., 2018). To investigate if varying substrate stiffness directly controls DLC1 expression, primary human umbilical vein endothelial cells (HUVECs) were cultured on fibronectin-coated 2 kPa (soft), 50 kPa (intermediate stiffness) and plastic (stiff) substrates. Western blot analysis of HUVEC lysates demonstrated elevated DLC1 protein levels on stiffer substrates (Fig. 1a). To study whether DLC1 upregulation in stiff microenvironments occurs through transcription regulation we performed quantitative PCR (QPCR) on RNA isolations from HUVECs. These experiments showed that mRNA levels of DLC1 are increased on stiff substrates (Fig. 1b). To investigate how DLC1 transcription is controlled by stiffness, we explored the promoter region of the DLC1 gene. We found a TEAD binding motif (CATTCCA) close to the transcriptional start site of the predominant DLC1 transcript variant expressed in endothelial cells (transcript variant 2, encoding for a 123 kDa protein). Analysis of publicly available TEAD1 chromatin immunoprecipitation sequencing data sets showed that TEAD1 binds this particular motif in a variety of different cell types (Fig. 1c and Suppl. Fig. 1). To study if the TEAD binding motif in the promoter region of DLC1 transcript variant 2 regulates promoter activity, wild-type or a TEAD binding motif-mutated variant of the promoter region of DLC1 (−418 to +319 bp) was fused to a luciferase reporter gene. Luciferase activity was monitored in lysates of transfected HEK cells cultured on plastic (stiff) substrates, which showed that mutating the TEAD motif perturbed transcriptional activation of the DLC1 promoter (Fig. 1d). Because YAP and TAZ act as co-factors for the TEAD family of transcription factors (Vassilev et al., 2001; Astone et al., 2018), we next investigated if YAP or TAZ are responsible for stiffness-induced DLC1 expression in endothelial cells. We performed short hairpin RNA (shRNA)-based knockdowns of YAP and TAZ in HUVECs cultured on plastic (stiff) substrates. Western blot analysis showed a strong reduction in the expression of DLC1 upon knockdown of YAP or TAZ (Fig. 1e and f), as well as the known YAP/TAZ-TEAD target connective tissue growth factor (CTGF) (Zhao et al., 2008).

**Figure 1.**
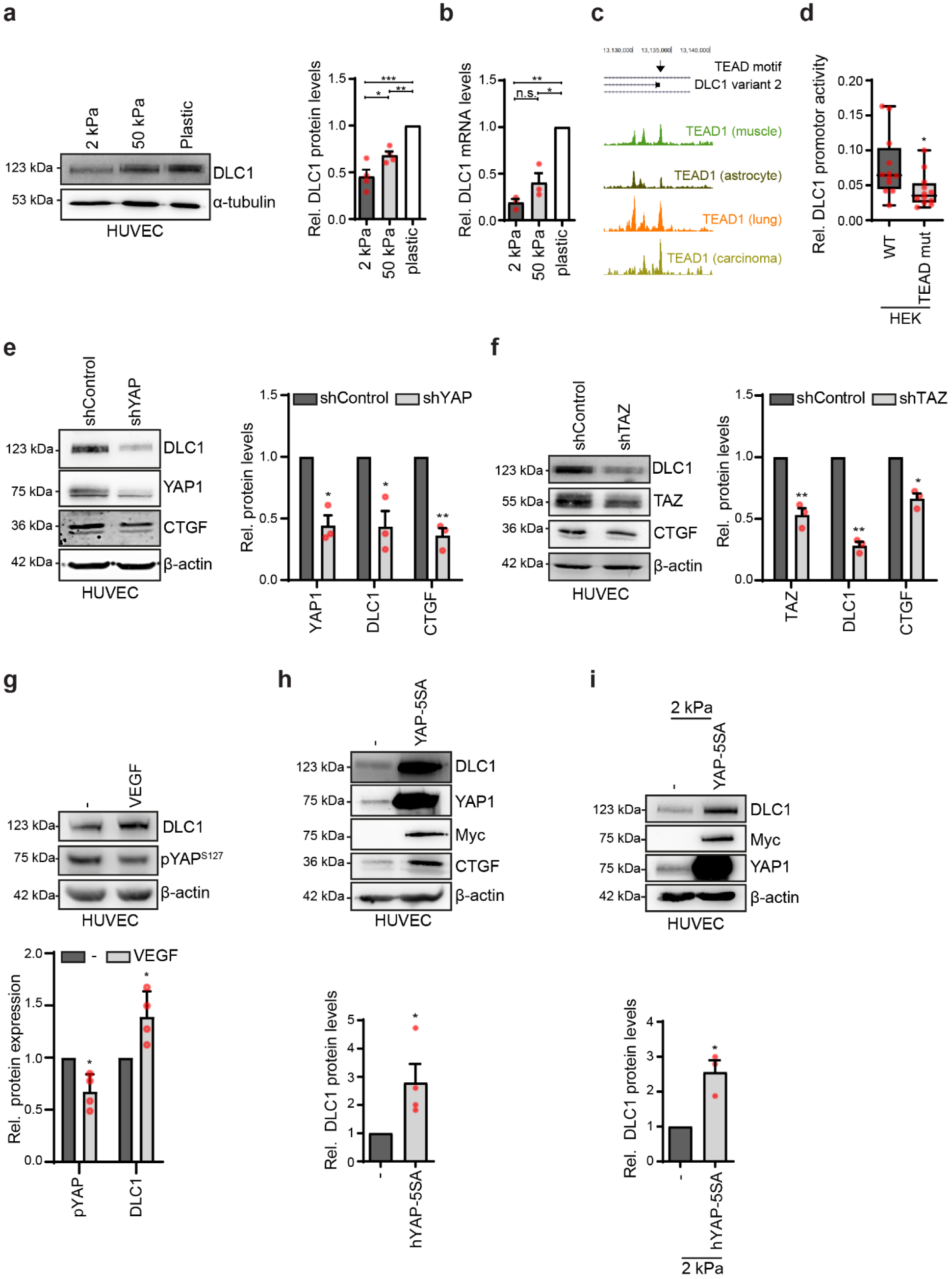
DLC1 is a transcriptional target of YAP/TAZ and TEAD. **(a)** Representative Western blot analysis of DLC1 and α-tubulin (loading control) in total lysates of HUVECs cultured on fibronectin-coated 2 kPa or 50 kPa hydrogels, or plastic. Graphs show the average protein expression levels ±s.e.m. Signal corrected for background and relative to expression in HUVECs on plastic. Data from 4 independent experiments. *P<0.05, **P<0.01, ***P<0.001 (One-way ANOVA and Dunnett’s post-test). **(b)** Quantitative real time PCR analysis of DLC1 mRNA isolated from HUVECs cultured on fibronectin-coated 2 kPa or 50 kPa hydrogels, or plastic. Graph shows the average gene expression levels ±s.e.m. Values are normalized to housekeeping gene (RPLP0) expression levels and relative to mRNA expression levels in HUVECs on plastic. Data from 3 independent experiments. n.s.; not significant, *P<0.05, **P<0.01 (One-way ANOVA and Dunnett’s post-test). **(c)** Schematics of UCSC genome browser results at position chr8:13,074,715-13,142,890 of the human genome (GRCh38/hg38) displaying the genomic location of DLC1 transcript variants 1 (NP872584.2), 2 (NP.006085.2) and 5 (NP.001303597.1) and the presence of a TEAD motif at the transcriptional start site of DLC1 transcript variant 2. Plotted are the results from publicly available GEO data TEAD1 ChIP-Seq data from various cell types, the data show a binding peak of TEAD1 at the TSS of DLC1 isoform 2. See Supplementary. Figure 1 for more details including histone modification profiles and DNase hypersensitivity profiles of the promoter region in HUVECs. **(d)** Boxplot showing the mean (± upper and lower quartiles) relative promoter activity of DLC1 in lysates of HEK cells transfected with wild type human DLC1 promoter region (from -418 to +319 bp position of the transcriptional start site of DLC1 isoform 2) fused to a firefly luciferase reporter or a DLC1-promoter luciferase reporter in which the TEAD-binding motif was mutated from CATTCCA to AGACTAT. Firefly luciferase activities were normalized to co-transfected renilla luciferase activity. Data is from 3 independent experiments. Whiskers show min/max values. *P<0.05 (Two-tailed unpaired Student’s t-test). **(e, f)** Representative Western blot analysis of DLC1 in total lysates of HUVECs transduced with shRNA Control, shRNA YAP **(e)** or shRNA TAZ **(f)**. Graphs show the average protein expression levels ±s.e.m. Signal corrected for background and normalized to expression in shControl-transduced HUVECs. Data from 3 independent experiments. *P <0.05, **P <0.01, ***P<0.001 (Paired Student’s t-test). **(g)** Representative Western blot analysis of DLC1, pYAP Serine 127 and β-actin (loading control) in total cell lysate samples of non-stimulated HUVECs (-) or HUVECs stimulated with 1mg/ml VEGF for 2 hours. Graphs show the average protein expression levels ±s.e.m. Signal corrected for background and normalized to expression in non-transduced HUVECs. Data from 4 independent experiments. *P<0.05 (Paired Student’s t-test). **(h)** Representative Western blot analysis of DLC1, YAP1, Myc, CTGF and β-actin (loading control) in total lysates of non-transduced HUVECs (-) and HUVECs transduced with Myc-tagged human YAP-5SA. Graphs show the average protein expression levels ±s.e.m. Signal corrected for background and normalized to expression in non-transduced HUVECs. Data from 4 independent experiments. *P<0.05 (Paired Student’s t-test). **(i)** Representative Western blot analysis of DLC1, YAP1, Myc, CTGF and β-actin (loading control) in total lysates of non-transduced HUVECs (-) and HUVECs transduced with Myc-tagged human YAP-5SA cultured on 2 kPa hydrogels. Graphs show the average protein expression levels ±s.e.m. Signal corrected for background and normalized to expression in non-transduced HUVECs. Data from 3 independent experiments. *P<0.05 (Paired Student’s t-test). Scans of whole Western blots are depicted in Supplemental Figure 2.

To establish whether activation of YAP/TAZ through other upstream cues might control DLC1 expression levels, serum and growth factor-starved HUVECs were treated with VEGF for 2 hours. Indeed, the VEGF treatments readily activated YAP, as analyzed by reduced phosphorylation of its Serine 127 as reported before (Wang et al., 2017), and upregulated expression levels of DLC1 (Fig. 1g). Next, a constitutive nuclear YAP-5SA mutant that cannot be inactivated by LATS1/2 kinases of the Hippo pathway (Zhao et al., 2007), was expressed in HUVECs. Expression of YAP-5SA strongly upregulated DLC1 and CTGF expression in HUVECs cultured on plastic substrates (Fig. 1h). To investigate if the constitutive active YAP-induced expression of DLC1 depends on substrate stiffness, control and YAP-5SA expressing HUVECs were cultured on 2 kPa substrates. Western blot analysis demonstrated that YAP-5SA efficiently promoted DLC1 expression even on soft substrates (Fig. 1i). Overall, these results demonstrate that DLC1 is a transcriptional target of YAP/TAZ and TEAD in the endothelium, and that YAP activation is sufficient to drive DLC1 expression.

### DLC1 controls endothelial focal adhesion turnover and traction forces

To investigate the role of DLC1 as a function of YAP/TAZ in the endothelium we silenced DLC1 expression through shRNAs. Three of the five tested shRNA plasmids induced efficient knockdown of DLC1 protein levels in HUVECs (Fig. 2a) and shDLC1 plasmids #1063 and/or #1064 were used for follow-up experiments. DLC1 may function at cadherin-based cell-cell junctions and integrin-based focal adhesions (Tripathi et al., 2012; Zacharchenko et al., 2016; Qian et al., 2007). DLC1 knockdown resulted in the formation of prominent basal F-actin fibers in endothelial cells, while the cells maintained their VE-cadherin-based cell-cell junctions (Fig. 2b). As YAP/TAZ are required for VE-cadherin dynamics and cell-cell junction formation in the vasculature (Neto et al., 2018), we first investigated the role of DLC1 in endothelial barrier function by Electric Cell-Substrate Impedance Sensing (ECIS). Upon knockdown of DLC1, no significant changes in endothelial barrier formation and maintenance of the barrier in time were detected (Fig. 2c). Since DLC1 is a GAP for Rho GTPases (Kim et al., 2009) we next compared the GTP-loading of RhoA in lysates of shControl and shDLC1 expressing HUVECs by means of G-LISA. We detected no differences in either basal or thrombin-stimulated RhoA activity levels (Fig. 2d). These data indicate that endothelial DLC1 is not required for endothelial cell-cell junctions, barrier function or RhoA activation.

**Figure 2.**
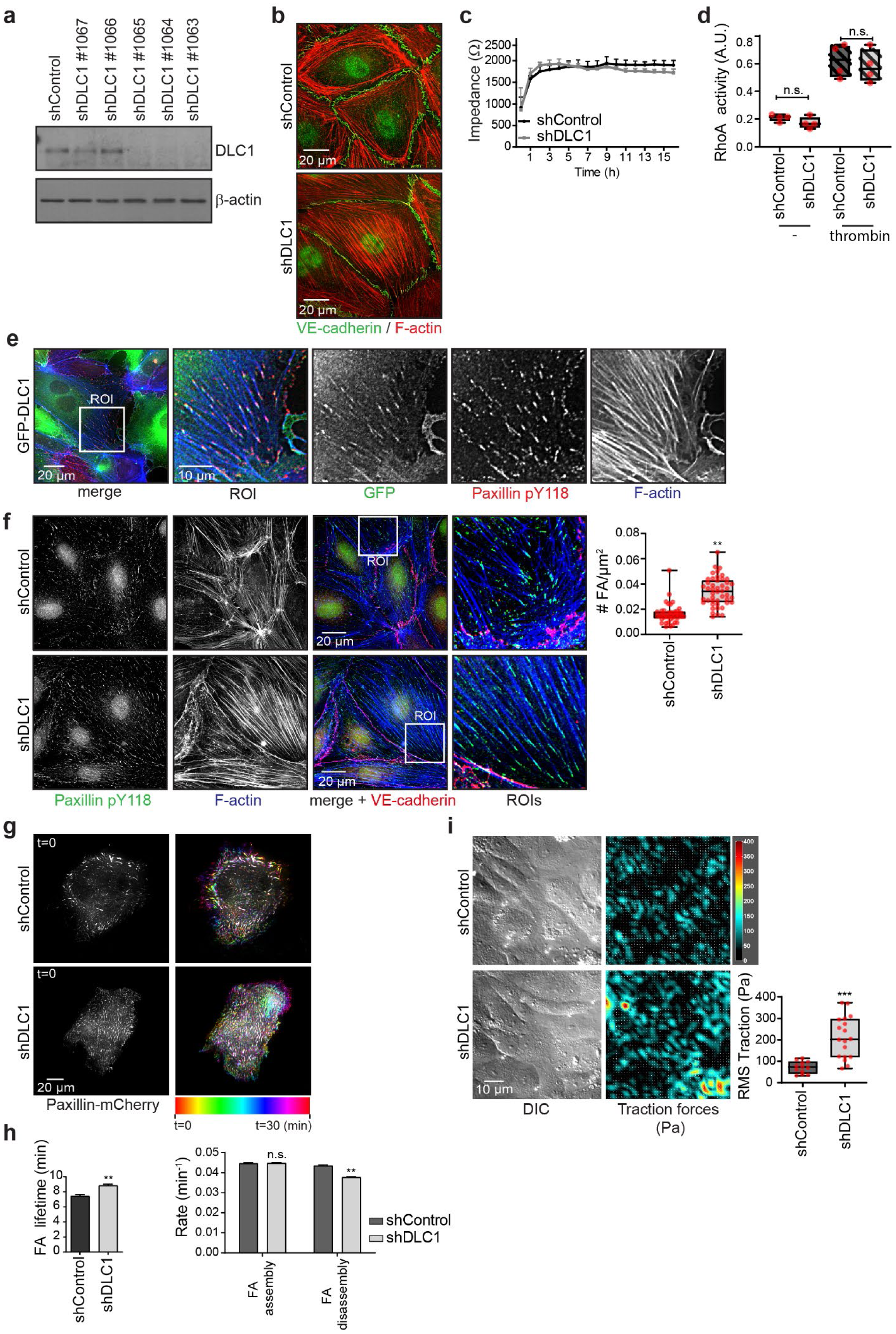
DLC1 controls endothelial focal adhesion turnover and traction forces. **(a)** Representative Western blot analysis of DLC1 and β-actin (loading control) in total lysates of HUVECs transduced with different shDLC1 clones (#1067, #1066, #1065, #1064, #1063). **(b)** Widefield immunofluorescence (IF) images of HUVECs transduced with shControl or shDLC1 (pool of #1063/1064) and stained for VE-cadherin (green) and F-actin (red). **(c)** Line graph shows the average ± stdev transendothelial impedance measured at 4000 Hz across barrier forming endothelial cells transduced with shControl or shDLC1 (pool of #1063/1064) plated on fibronectin-coated 8W10E ECIS arrays. Representative data is from 2 independent experiments and an average of 6 wells per condition. **(d)** Boxplot showing quantification of RhoA activity in G-LISA assays in lysates of shControl and shDLC1 transduced HUVECs, with or without thrombin stimulation. Data is from 4 independent experiments. Whiskers show min/max values. n.s.; not significant (Two-tailed unpaired Student’s t-test). **(e)** Representative widefield IF images of HUVECs transduced with GFP-DLC1 stained for Paxillin pY118 (red) and F-actin (blue). **(f)** Widefield IF images of shControl and shDLC1 transduced HUVECs and stained for Paxillin pY118 (green), F-actin (blue), VE-cadherin (red). Boxplot showing quantification of manually counted number of focal adhesions per µm2 of shControl and shDLC1 transduced HUVECs. Whiskers show min/max values. Data is from 2 independent experiments, shControl (42 cells from 12 images) and shDLC1 (46 cells from 19 images). **P<0.01 (Two-tailed unpaired Student’s t-test). **(g)** Left images are stills from time lapse TIRF microscopy at t=0 of HUVECs transduced with shControl and shDLC1 and paxillin-mCherry. Heat map shows the corresponding focal adhesion dynamics over 30 minutes in a unique color per time frame. Note the stability of focal adhesions in shDLC1 HUVECs as indicated by the white colored FAs in the right panels. See corresponding Supplemental Movie 1 for the ∼2.5 hours time-lapse recording. **(h)** Bar graph showing quantification of focal adhesion lifetime, assembly and disassembly rates based on TIRF time lapse experiments with paxillin-mCherry expressing HUVECs. Error bars are s.e.m. Data is from 2 independent experiments, shControl (6 movies) and shDLC1 (8 movies). Focal adhesion tracking was performed using the focal adhesion analysis webserver (Berginski and Gomez, 2013). **P<0.01 (Two-tailed unpaired Student’s t-test). **(i)** DIC images and cell-substrate traction force maps of HUVECs transduced with shControl or shDLC1. Boxplot showing the mean (± upper and lower quartiles) of measured RMS traction forces of shControl (12 image fields) and shDLC1 (18 image fields) endothelial cells. Data is from 3 independent experiments. Whiskers show min/max values. **P<0.01 (Two-tailed unpaired Student’s t-test).

Lentiviral expression of an N-terminal GFP-tagged DLC1 (Qian et al., 2007) showed that DLC1 is recruited to focal adhesions in HUVECs (Fig. 2e). Next, we investigated the role of DLC1 at endothelial integrin-based adhesions by immunofluorescence staining for paxillin in shControl and shDLC1 expressing HUVECs. These experiments showed that depletion of DLC1 strongly increased the number of focal adhesions that were connected to their prominent F-actin fibers (Fig. 2f). Integrin-based focal adhesions are highly dynamic structures, that are constantly being formed and disassembled (Möhl et al., 2012; Geiger et al., 2009; Gardel et al., 2010). To decipher the mechanism underlying the remodeling of focal adhesions by DLC1, we performed live imaging using total internal reflection fluorescence microscopy (TIRF) of shControl and shDLC1 HUVECs expressing mCherry-tagged paxillin. Consistent with our previous findings, the TIRF imaging showed that DLC1-depleted HUVECs contained more focal adhesions (Fig. 2g and Suppl. Movie 1; note only ∼10-20% of the cell population in the monolayer is paxillin-mCherry positive). Overall, the change in adhesion turnover in the absence of DLC1 resulted in a striking stabilization of the focal adhesions (Fig. 2g, white colored focal adhesions in right panels). Quantitative analysis using established focal adhesion tracking software (Berginski and Gomez, 2013) showed an increase in focal adhesion lifetime, which corresponded with a decrease in focal adhesion disassembly rates, while assembly rates remained comparable (Fig. 2h). To investigate the functional consequences of depletion of DLC1 for force transduction from cells to the ECM, traction force microscopy (TFM) was performed using shControl and shDLC1 expressing HUVECs. These experiments demonstrated that depletion of DLC1 promoted endothelial traction forces throughout the monolayer, and strongly raised the root mean square of exerted traction forces (Fig. 2i). Thus, DLC1 expression is driven by substrate stiffness, and in turn, expression of DLC1 controls force transduction at the cell-ECM interface. Together, the data show that DLC1 is needed for efficient turnover of endothelial focal adhesions and traction forces.

### Endothelial DLC1 controls cell orientation and directed migration

In cell collectives YAP/TAZ translocate to the nuclei of leader cells to regulate endothelial cell migration (Lin et al., 2017; Yu and Guan, 2013; Zhang et al., 2015; Mason et al., 2019; Neto et al., 2018)(and Suppl. Fig. 3). We next examined if endothelial DLC1 is involved in collective cell migration. Knockdown of DLC1 strongly inhibited endothelial migration in scratch wound assays (Fig. 3a, b and Suppl. Movie 2), supporting previous findings for a role of DLC1 in migration of prostate epithelial cells (Shih et al., 2012). Within 12 hours, control monolayers closed on average 88.89% of the wound area, while DLC1 knockdown inhibited scratch wound closure: 41.66% and 67.16% for an shRNA targeting the 3’ UTR of DLC1 mRNA and clone #1063 respectively (Fig. 3b, c). No changes in cell proliferation were observed between the conditions (Fig. 3d), indicating that the delay in wound healing was due to migration defects. To investigate how DLC1 controls cell dynamics, we performed tracking of individual endothelial cells within the confluent monolayers in time lapse experiments. Whereas control cells migrated collectively in a persistent fashion in the direction of wound closure, DLC1-depleted cells lost their capability for directional migration (Fig. 3e and Suppl. Movie 2). These data demonstrate that DLC1 is needed to coordinate collective cell migration.

**Figure 3.**
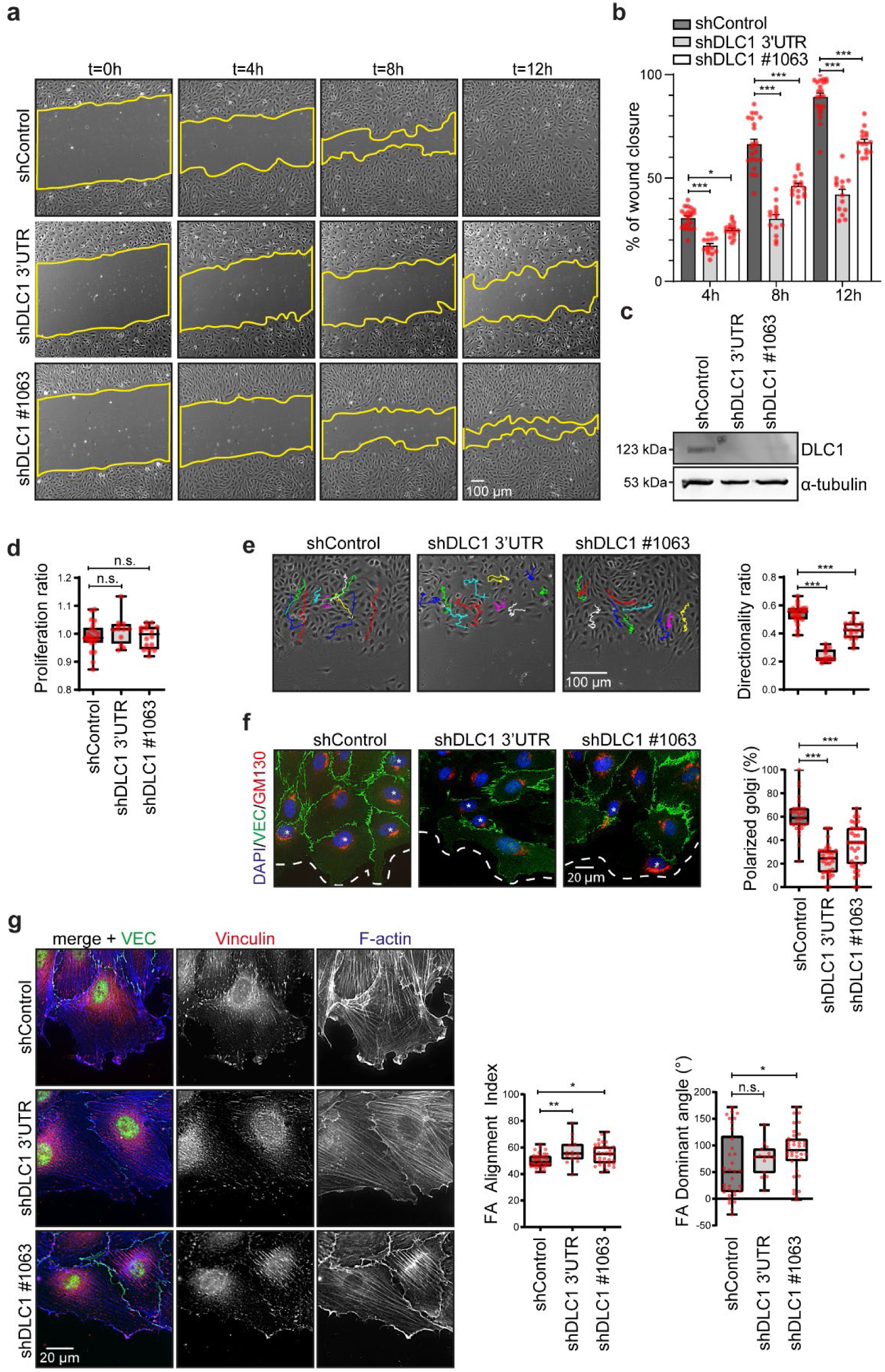
DLC1 is required for cell orientation and directional migration. **(a)** Representative Phase-contrast images of HUVECs transduced with shControl, shDLC1 3’UTR or shDLC1 #1063, in a scratch-wound assay (t=0, t=4, t=8 and t=12 hours after scratch). Yellow line highlights the unclosed wound area. See corresponding Supplemental Movie 2 for the ∼16 hours timelapse recording. **(b)** Graphs showing the percentage of wound closure of HUVECs transduced with shControl, shDLC1 3’UTR or shDLC1 #1063 at three time points during scratch-wound assay. Data is from 4 independent experiments, shControl (22 movies), shDLC1 3’UTR (14 movies) and shDLC1 #1063 (18 movies). *P<0.05, ***P<0.001 (Two-way ANOVA and Tukey’s multiple comparison test). **(c)** Representative Western blot analysis of DLC1 and α-tubulin (loading control) in total lysates of HUVECs transduced with shControl, shDLC1 3’UTR or shDLC1 #1063. Scans of whole Western blots are depicted in Supplemental Figure 4. **(d)** Boxplot showing the mean (± upper and lower quartiles) of proliferation ratio of the shControl, shDLC1 3’UTR, shDLC1 #1063 HUVECs corresponding to the scratch assays in Fig. 3b. Proliferation ratio was manually determined by counting cell numbers at ROIs within the endothelial monolayers before and after 12 hrs scratch wound migration. n.s.; not significant (One-way ANOVA with Dunnett’s multiple comparison test). **(e)** Phase-contrast images of scratch wound assays overlayed with single cell tracking analysis of 8 hours using the Chemotaxis Tool in ImageJ. Boxplot showing the mean (± upper and lower quartiles) velocity and directionality of shControl, shDLC1 3’UTR or shDLC1 #1063 transduced HUVECs. Whiskers show min/max values. Data is from 4 independent experiments, shControl (429 cells from 22 movies), shDLC1 3’UTR (236 cells from 12 movies), shDLC1 #1063 (303 cells from 18 movies). n.s.; not significant, ***P<0.001 (One-way ANOVA and Dunnett’s multiple comparison test). **(f)** Widefield IF images of HUVECs transduced with shControl, shDLC1 3’UTR or shDLC1 #1063 6 hours after initiation of scratch wound assays stained for DAPI (blue), GM130 (red; Golgi) and VE-cadherin (green). The dotted lines indicate migration front and the asterisks indicate cells containing an oriented Golgi in the direction of scratch wound migration. Boxplot showing the mean (± upper and lower quartiles) of cells with an oriented Golgi. Data is from 3 independent experiments, shControl (34 images), shDLC1 3’UTR (30 images), shDLC1 #1063 (35 images). ***P<0.001 (One-way ANOVA with Dunnett’s multiple comparison test). **(g)** Widefield IF images of HUVECs transduced with shControl, shDLC1 3’UTR or shDLC1 #1063 6 hours after initiation of scratch wound assays stained for Vinculin (red), F-actin (blue) and VE-cadherin (green). Boxplots show the mean (± upper and lower quartiles) Focal Adhesion Alignment Index and Focal Adhesion dominant angle as determined by image analysis of the vinculin channels using the Focal Adhesion Analysis Server (Berginski and Gomez, 2013). Data is from 3 independent experiments, shControl (34 images), shDLC1 3’UTR (15 images), shDLC1 #1063 (36 images). Whiskers show min/max value. n.s.; not significant, *P< 0.05, **P<0.01 (One-way ANOVA with Dunnett’s multiple comparison test).

Cell polarization is needed for persistent migration, which is characterized by the orientation of the Golgi apparatus in front of the nucleus (Kupfer et al., 1982; Bisel et al., 2013). To study if DLC1 controls cell polarization during migration, shControl and shDLC1 expressing endothelial monolayers were analyzed 6 hours after the initiation of scratch wound migration. Golgi orientation was determined in the first three leader cell rows in immunostainings for GM130 (Golgi marker) and considered polarized if located within an angle between 60° and -60° towards the migration front. In control HUVECs, 60% of the cells oriented their Golgi in the direction of migration, whereas only 23% to 37% of DLC1-depleted cells were polarized (Fig. 3f). To investigate whether the failure of DLC1 knockdown cells to polarize relates to differences in focal adhesion turnover, we investigated the alignment of focal adhesions in leader cells 6 hours after the onset of migration. Immunostainings for vinculin, a marker of focal adhesions, showed that in the absence of DLC1 the focal adhesions aligned more among each other than in control cells, but the aligned focal adhesions oriented perpendicularly (>90° dominant angle) to the direction of scratch wound closure (Fig. 3g). Overall, these results clearly show that DLC1 is needed for endothelial cell polarization and focal adhesion organization during collective cell migration.

### DLC1 is required for sprouting angiogenesis

Directional migration is essential for endothelial remodeling during sprouting angiogenesis (Laurent et al., 2007; Eilken and Adams, 2010; Franco et al., 2015). Moreover, the sensing of ECM stiffness and exertion of tensional forces occurs through endothelial integrin-based adhesions and directs the formation of angiogenic sprouts (Fischer et al., 2018; Korff and Augustin, 1999). To establish whether DLC1 plays a role in angiogenic sprouting, HUVECs expressing shControl and shDLC1 were cultured in spheroids and placed in 3D collagen matrices to assess sprouting capacity as described previously (Heiss et al., 2015; Martin et al., 2018). Subsequently, sprouting was induced by treatment with VEGF. Visualization of sprout formation after 16 hours, showed a decrease in cumulative length and the number of sprouts after depletion of DLC1 (Fig. 4a). Moreover, overexpression of GFP-DLC1 efficiently promoted sprout formation compared to GFP-transduced control HUVECs (Fig. 4b). These data indicate that expression levels of DLC1 determine angiogenic sprouting efficiency. To verify the contribution of DLC1 in sprouting angiogenesis, we first depleted endogenous DLC1 by shRNAs targeting the 3’ UTR of the mRNA. Subsequently, GFP or GFP-DLC1, that are not targeted by the shRNAs, were expressed. Restoring expression of DLC1 efficiently rescued endothelial sprouting capacity (Fig. 4c). Western blot analysis confirmed the knockdown and expression of the GFP-tagged DLC1 (Fig. 4d). Overall, the data indicate that endothelial DLC1 expression levels tightly control sprouting angiogenesis.

**Figure 4.**
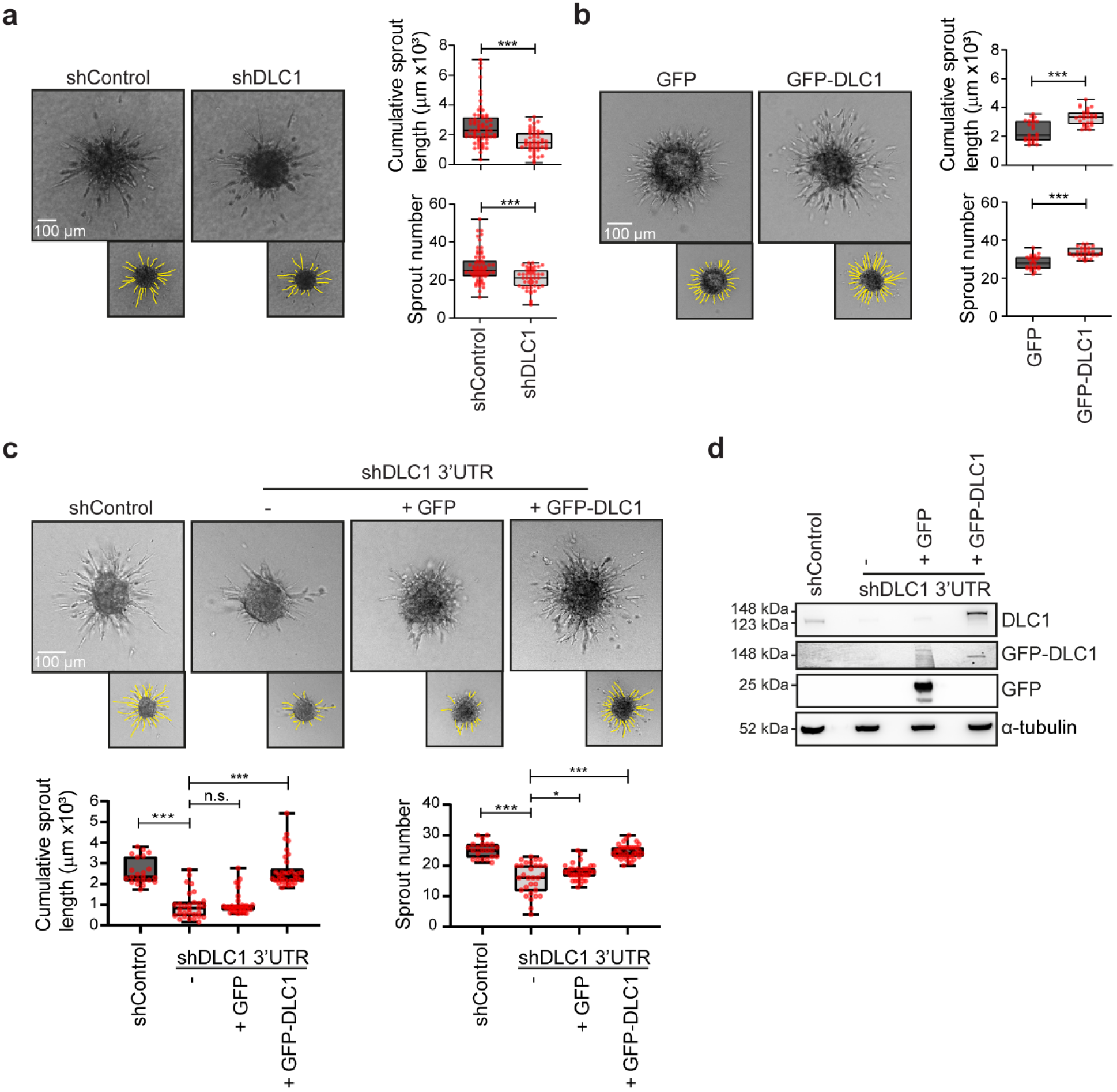
DLC1 controls sprouting angiogenesis. **(a)** Representative phase-contrast images of sprouting spheroids 16 hours after VEGF stimulation of HUVECs lentivirally transduced with shControl, shDLC1 (#1064/1065). Boxplot shows the mean (± upper and lower quartiles) cumulative sprout length and the number of sprouts in the spheroid-based sprouting angiogenesis assays. Whiskers show min/max values. Data is from 3 independent experiments, shControl (63 spheroids) and shDLC1 (47 spheroids). **P<0.01 (Two-tailed unpaired Student’s t-test). **(b)** Representative phase-contrast images of sprouting spheroids 16 hours after VEGF stimulation of HUVECs lentivirally transduced with GFP or GFP-DLC1. Boxplot shows the mean (± upper and lower quartiles) cumulative sprout length and the number of sprouts in the spheroid-based sprouting angiogenesis assays. Whiskers show min/max values. Data is from 3 independent experiments, GFP (25 spheroids), GFP-DLC1 (27 spheroids). ***P<0.001 (Two-tailed unpaired Student’s t-test). **(c)** Representative phasecontrast images of sprouting spheroids 16 hours after VEGF stimulation of HUVECs transduced with shControl or shDLC1 3’UTR and rescued with GFP, GFP-DLC1 or GFP-DLC1-R677E. Boxplots show the mean (± upper and lower quartiles) cumulative sprout length and number of sprouts per spheroid. Whiskers present min/max values. Data is from 3 independent experiments, shControl (33 spheroids), shDLC1 3’UTR (29 spheroids), rescue GFP (39 spheroids), rescue GFP-DLC1 (37 spheroids), rescue GFP-DLC1 R667E (28 spheroids). n.s. non-significant, *P<0.05, ***P<0.001 (One-way ANOVA with Dunnett’s multiple comparisons test). **(d)** Representative Western blot analysis of DLC1, GFP and α-tubulin (loading control) in lysates of HUVECs transduced with shControl or shDLC1 3’UTR and rescued with GFP or GFP-DLC1. Note that the GFP-tagged DLC1 has a higher molecular weight than endogenous DLC1. Scans of whole Western blots are depicted in Supplemental Figure 4.

### DLC1 rescues the migration and sprouting defects in YAP-depleted endothelial cells

Endothelial YAP/TAZ activation drives angiogenesis by controlling endothelial collective migration and vessel remodeling (Neto et al., 2018; Kim et al., 2017). The transcriptional targets of YAP/TAZ that are responsible for this task remain unknown. Since DLC1 is needed for collective migration and angiogenic sprouting, we next assessed the contribution of DLC1 as downstream target of YAP. First, YAP expression was silenced using shRNAs. The knockdown of YAP in HUVECs, which is accompanied by a downregulation of DLC1 expression, inhibited scratch wound migration (Fig. 5a, b, c and 1e), confirming previous findings (Neto et al., 2018). IF imaging further revealed that the defective scratch wound migration of YAP-depleted HUVECs is accompanied by the formation of perpendicularly oriented focal adhesions and actin stress fibers in cells at the leading edge (Fig. 5d), reminiscent of the morphology of DLC1-depleted HUVECs in scratch wound assays. To investigate the contribution of DLC1 as target of YAP in focal adhesion remodeling during collective migration, DLC1 protein levels were restored in shYAP HUVECs by ectopic expression of GFP-DLC1 (Fig. 5c). Ectopic expression of DLC1 in shYAP HUVECs, induced proper alignment of focal adhesions and the actin cytoskeleton of cells at the leading edge and partially rescued the collective cell migration defects of YAP-depleted cells (Fig. 5a, b, c, d). Next, to address the contribution of DLC1 in YAP-dependent sprouting angiogenesis, the cells were analyzed for their sprouting capacity. Intriguingly, restoring DLC1 levels by ectopic expression of DLC1 fully rescued the sprouting defects of YAP-depleted HUVECs (Fig. 5e). Taken together, these findings point towards a crucial and prominent role for DLC1 in YAP/TAZ-driven endothelial adhesion remodeling and collective migration during angiogenesis.

**Figure 5.**
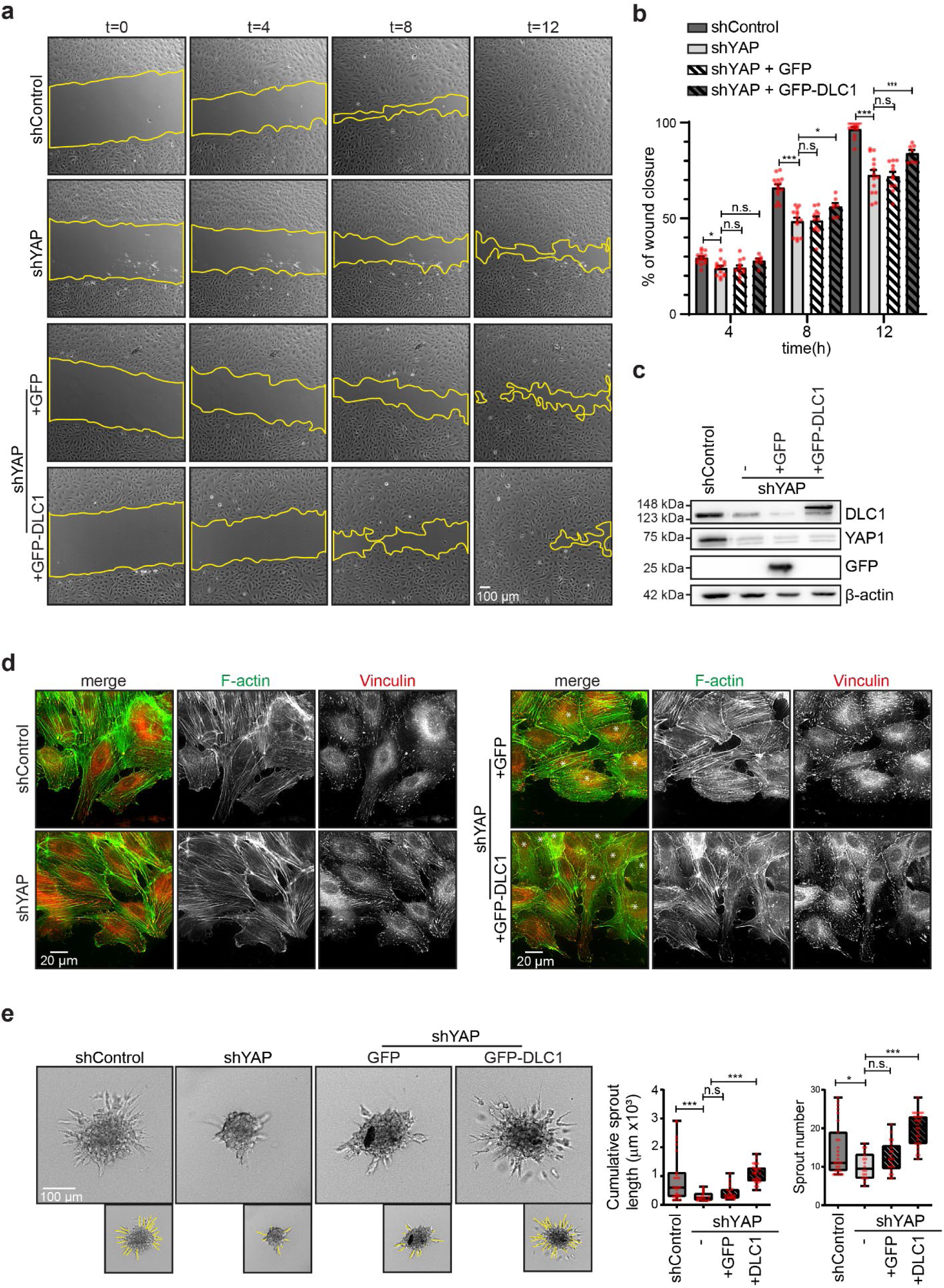
DLC1 rescues the migration and sprouting defects in YAP-depleted endothelial cells. Phase-contrast images of HUVECs transduced with shControl or shYAP1 and rescued with GFP or GFP-DLC1, in a scratch-wound assay (t=0, t=4, t=8 and t=12 hours after scratch). Yellow line highlights the unclosed wound area. Graphs showing the percentage of wound closure of HUVECs transduced with shControl or shYAP and rescued with GFP or GFP-DLC1 at three time-points during scratch-wound assay. Data is from 3 independent experiments, shControl (14 movies), shYAP (14 movies), rescue GFP (11 movies), rescue GFP-DLC1 (7 movies). n.s. nonsignificant, *P<0.05, ***P<0.001 (One-way ANOVA with Dunnett’s multiple comparisons test). **(c)** Western blot analysis of DLC1, YAP, GFP and □-actin in lysates of HUVECs transduced with shControl, shYAP and rescued with GFP or GFP-DLC1. Scans of whole Western blots are depicted in Supplemental Figure 4. **(d)** Representative widefield IF images of HUVECs transduced with shControl or shYAP, or shYAP-transduced HUVECs expressing GFP or GFP-DLC1, 6 hours after initiation of scratch wound assay. Stained for Vinculin (red) and F-actin (green). The asterisks indicate the GFP positive cells (not shown). **(e)** Representative phasecontrast images of sprouting spheroids 16 hours after VEGF stimulation of HUVECs transduced with shControl, shYAP and rescued with GFP or GFP-DLC1. Boxplots show the mean (± upper and lower quartiles) cumulative sprout length and number of sprouts per spheroid. Whiskers present min/max values. Data is from 3 independent experiments, shControl (27 spheroids), shYAP (18 spheroids), rescue GFP (21 spheroids), rescue GFP-DLC1 (28 spheroids). n.s.; not significant, *P<0.05, ***P<0.001 (One-way ANOVA with Dunnett’s multiple comparison test).

## Discussion

The nuclear translocation of YAP/TAZ during stiffness and flow sensing tightly controls cell-ECM interactions, yet the downstream targets of YAP/TAZ that are responsible for such mechano-responses still remain unclear (Totaro et al., 2018). Our study reveals that the focal adhesion protein DLC1 is a direct transcriptional target of YAP/TAZ and TEAD and crucial for YAP-driven collective cell migration and sprouting angiogenesis by endothelial cells. These findings implicate DLC1 in related YAP/TAZ-driven mechanotransduction processes, such as flow sensing, contact inhibition and the development of stiffness-related vascular disease.

### DLC1 and regulation of endothelial adhesion dynamics

YAP/TAZ are important mechanotransducers that translocate to the nucleus upon cell-ECM adhesion-induced actomyosin tension (Dupont et al., 2011). In turn YAP/TAZ activation has been shown to control focal adhesions in various cell types (Nardone et al., 2017). Endothelial focal adhesions have recently been shown to provide feedback signals to YAP/TAZ activity to limit adhesion maturation for cell orientation and persistent migration (Mason et al., 2019). Intriguingly, our data now demonstrate that DLC1, following activation of YAP/TAZ, restricts focal adhesion lifetime, confines integrin-based traction forces and thereby promotes cell polarization during directed migration. These findings suggest that upregulation of DLC1 expression upon YAP/TAZ activation provides the feedback signals for optimal adhesion remodeling and force transduction.

It is still uncertain how DLC1 controls the turnover of focal adhesions and endothelial dynamics. Adhesion turnover is steered by spatiotemporal activation of Rho GTPases and subsequent cytoskeletal remodeling (Webb et al., 2002; Etienne-Manneville and Hall, 2002). DLC1 is widely known as a Rho GAP protein to inhibit GTP-loading of Rho GTPases (Wong et al., 2003; Healy et al., 2008). The GAP activity of DLC1 contributes to the tumor suppressive functions of DLC1 (Healy et al., 2008; Ko et al., 2013). However, we find no major differences in RhoA activation upon depletion of DLC1, pointing towards an alternative function of DLC1 in endothelial adhesion remodeling. In epithelial cells DLC1 is involved in migration through a GAP-independent manner (Shih et al., 2012), and in HeLa cells the recruitment of DLC1 to focal adhesions is needed for its pro-migratory function (Kawai et al., 2009). DLC1 has been shown to interact with the integrin-related proteins Talin, focal adhesion kinase (FAK) and Tensin (Qian et al., 2007; Liao and Lo, 2008; Li et al., 2011; Zacharchenko et al., 2016). Interestingly, tension-induced conformational unfolding of Talin, a key mechanosensor for integrins, inhibits the interaction between DLC1 and Talin and prevents downstream inhibition of myosin phosphorylation (Haining et al., 2018). Thus DLC1’s interaction with Talin might regulate myosin-driven turnover of focal adhesions to favor migration. Of note, in fibroblasts and various cancer cell types DLC1 seems to have an opposing role in cell migration (Heering et al., 2009; Barras and Widmann, 2014; Kaushik et al., 2014), which might relate to differences in YAP/TAZ activation and indicates that DLC1’s function is highly dependent on the cellular microenvironment. Within blood vessels, mechanotransduction through integrins predominantly occurs in arterial endothelium (Van Geemen et al., 2014; Di Russo et al., 2017), suggesting that the regulation of focal adhesions by DLC1 could be of particular importance in arteries.

### The role of YAP/TAZ and DLC1 in angiogenesis

YAP/TAZ signaling serves prominent roles in vascular biology and angiogenesis (Choi et al., 2015; Nakajima et al., 2017a; Wang et al., 2016a, 2017; Neto et al., 2018). YAP/TAZ is regulated by ECM rigidity, (blood) flow and cell density, mechanical cues which also influence angiogenesis (Zhao et al., 2007; Aragona et al., 2013; Wang et al., 2016b; a; Boerckel et al., 2011). In addition, the angiogenesis process is supported by mechanical stretch- and VEGF-mediated activation of YAP/TAZ (Neto et al., 2018; Wang et al., 2017). Interestingly, microarray analysis of HUVECs expressing constitutive nuclear YAP and TAZ mutants, which induce hypersprouting in angiogenesis, also resulted in an upregulation of DLC1 expression amongst the regulation of other transcriptional programs (Neto et al., 2018). We now reveal that DLC1 is needed for VEGF-induced sprouting and that ectopic DLC1 expression is sufficient to restore sprouting in YAP-depleted endothelial cells. DLC1 and the related protein DLC2 (also known as STARD13) have been shown to contribute to experimentally-induced angiogenesis in vivo (Shih et al., 2017; Lin et al., 2010). Moreover, DLC1 was described to regulate contact inhibition of growth in endothelial cells (Sánchez-Martín et al., 2018). This fits with a model in which DLC1 expression levels are tightly controlled for endothelial rearrangements during angiogenesis. The adherens junction receptor VE-cadherin, an endothelial specific cadherin that safeguards vascular integrity and steers endothelial dynamics (Carmeliet et al., 1999; Bentley et al., 2014), sequesters YAP at stabilized endothelial cell-cell junctions to prevent its activation (Giampietro et al., 2015). Possibly, the inhibition of YAP/TAZ nuclear translocation by VE-cadherin contributes to stabilization of blood vessel integrity through suppression of DLC1 driven-endothelial migration and/or angiogenic sprouting.

### Disturbed cross talk between DLC1 and YAP/TAZ in pathology?

DLC1 expression is elevated in the endothelium of atherosclerotic plaques and in pulmonary hypertension (Schimmel et al., 2018). The development of these cardiovascular diseases is driven by pathological stiffening and disturbed flow patterns (Huveneers et al., 2015), which activate endothelial YAP/TAZ (Wang et al., 2016a; b; Bertero et al., 2016). Therefore targeting of YAP/TAZ, or their downstream targets such as DLC1, holds promise as therapeutic approach in stiffness-related vascular diseases. Of interest, mutations in the YAP and TAZ genes (Antonescu, 2014) correlate with loss of DLC1 expression in hemangioendotheliomas and angiosarcomas (Sánchez-Martín et al., 2018; Ha et al., 2018), two (rare) types of endothelial malignancies that are characterized by uncontrolled proliferation and infiltration of endothelial cells. The latter reports also suggest that DLC1 deficiency is an upstream inducer of YAP/TAZ signaling in endothelial malignancies. This would fit with a model in which YAP/TAZ and DLC1 control each other via feedback mechanisms that are sensed at focal adhesions. Because DLC1 is expressed in many cell types beyond the endothelium, and because DLC1 has been identified as a tumor suppressor in various types of cancer (Liao and Lo, 2008; Durkin et al., 2007), we postulate that DLC1 might also function in YAP/TAZ-driven cancer cell behavior, and other epithelial tissue processes that involve remodeling of integrin-based adhesions.

## Materials & Methods

### Antibodies and reagents

Purified mouse anti-human DLC1 (Clone 3, Cat # 612021, diluted 1:1000 for Western) and purified mouse anti-GM130 (clone 35, Cat # 610823, diluted 1/200 for IF) were obtained from BD biosciences. To visualize VE-cadherin we used purified mouse anti-cadherin-5 (BD biosciences, clone 75, Cat # 610252, diluted 1/100 for IF) and rabbit polyclonal anti-VE-cadherin (Cayman Chemical, Cat # 160840, diluted 1/100 for IF). Rabbit polyclonal antiphospho-paxillin (Tyr118) (Cat # 44-722G, diluted 1/200 for IF) was from Thermo Fischer Scientific. Mouse monoclonal anti-vinculin (hVIN-1 clone, Cat # V9131, diluted 1/400 for IF) and rabbit polyclonal anti-WWTR1 (anti-TAZ, cat # HPA007415, diluted 1/1000 for western) were obtained from Sigma-Aldrich. We used rabbit polyclonal anti-YAP1 (Genetex, Cat # GTX129151, diluted 1/1000 for western) to detect YAP, mouse monoclonal anti-c-Myc (clone 9E10, Cat # SC-40, diluted 1/1000 for western) and goat polyclonal anti CTGF (Santa Cruz, L-20, Cat # sc-14939, diluted 1/1000 for western) to detect CTGF. To detect GFP we used the monoclonal mouse anti-GFP antibody (Santa Cruz, B-2, Cat # sc-9996, 1/1000 for western). As loading control we used mouse monoclonal anti-human α-Tubulin (Cedarlane, Clone DM1A, Cat # CLT9002, diluted 1/10.000 for western) or rabbit polyclonal anti-β-Actin (Cell Signaling, Cat # 4967S, diluted 1/1000 for western). Secondary antibodies coupled to Alexa Fluor-488, - 594, and -647 were purchased from Invitrogen (diluted 1/100 for IF). To visualize F-actin we used PromoFluor-415 Phalloidin (Promokine, Cat # PK-PF415-7-01, diluted 1/200 for IF), Alexa Fluor 568 Phalloidin (Thermo Fisher, Cat # A12380, diluted 1/200 for IF) or Texas Red-X Phalloidin (Thermo Fisher, Cat # T7471, diluted 1/200 for IF). DAPI was used for nuclear IF stainings (Invitrogen, diluted 1:1000). Secondary antibodies coupled to horseradish peroxidase (HRP) were obtained from Bio-Rad (Diluted 1/1000 for western). Thrombin (used at 0.2U/ml) was from Haematologic Technologies. Doxycyclin (used at 1 ng/ml) was from Sigma.

### Cell culture

Pooled primary HUVECs (cultured up to passage six) from different donors (Lonza) were cultured in Endothelial Cell Growth Medium 2 culture medium supplemented with the Growth Medium 2 Supplement Pack (PromoCell) on gelatin-coated tissue flasks. 2 kPa or 50 kPa hydrogels (Matrigen) were activated with PBS and coated with 5 µg ml-1 fibronectin overnight. HEK293T cells (ATCC) were cultured in Dulbecco’s Medium Eagle medium with L-glutamine supplemented with 10% fetal bovine serum and antibiotics.

### DNA plasmids and lentiviral transductions

shRNAs in the lentiviral pLKO.1 backbone targeting DLC1 (TRCN47823, 47824, 47825, 47826 and 47827, which in this manuscript are referred to plasmid numbers #1063, #1064, #1065, #1066 and #1067 respectively), YAP1 (TRCN107265), TAZ (TRCN19473) and control shRNA (shC002) were from Sigma Aldrich mission library. A modified version of the pLKO.1 plasmid was generated based on the GGAGTGTAGGAATTGACTATA sequence to express shRNA that target the 3’-UTR of human DLC1 mRNA. Full-length human DLC-1 fused at its N terminus to a GFP tag was amplified by PCR from a pEGFP-C1-DLC-1 vector (provided by Xiaolan Qian and Douglas Lowy) and cloned into a self-inactivating lentiviral pLV-CMV-ires-puro vector between the SnaBI and XbaI restriction sites. For the pInducer20-myc-hYAP1-5SA-Ubc construct, human YAP1 with S61A, S109A, S127A, S164A, S381A mutations was amplified by PCR from a pQCXIH vector (Zhao et al., 2007) and through Gateway Recombination cloned into a pInducer20 vector with Ubc promoter (Meerbrey et al., 2011). The pRRL paxillin-mCherry construct was a gift from Olivier Pertz (University of Basel, Switzerland). Luciferase reporter constructs are based on a pGL3 basic luciferase reporter vector (Promega), containing a Firefly Luciferase gene. The DLC1 promoter region (−418 to +319 bp from the transcriptional start site) containing the wild type TEAD motif (CATTCCA) or a mutated motif (AGACTAT) were purchased from GenScript and inserted at the 5’ end of the luciferase gene between NheI and BglII restriction sites. To produce lentiviral particles HEK293 cells were transfected with third-generation packaging constructs and lentiviral expression vectors using Trans-IT-LTI transfection reagents (Mirus). Lentivirus containing supernatant was harvested 48-72 hours post transfection. HUVECs, cultured up to ∼80% confluency, were transduced with lentiviral particles overnight. shRNA-based knockdown cells were analyzed at least 48 hours after transduction. For TIRF microscopy HUVECs were first transduced with shDLC1 or shControl lentivirus, and subsequently transduced with paxillin-mCherry lentivirus. Expression of YAP-5SA was induced by doxycyclin treatment for 48 hours.

### Electric cell-substrate impedance sensing (ECIS)

Electric cell-substrate impedance sensing was used to analyze endothelial barrier function. Electrode arrays (8W10E; IBIDI) were treated with 10 mM L-cysteine (Sigma) for 15 minutes at 37 °C. After washing with 0,9% NaCl, the arrays were coated with 10 µg ml-1 fibronectin in 0,9% NaCl for 1 hour at 37°C. Cells were seeded on the arrays and the impedance was measured during monolayer formation at 4 kHz using the ECIS model ZTheta (Applied BioPhysics).

### G-LISA - RhoA activity assays

For analysis of RhoA activity, confluent HUVECs were washed with ice-cold PBS and lysed in lysis buffer from the RhoA G-LISA activation kit (Cytoskeleton). RhoA activity was determined according to manufacturer’s protocol.

### Immunofluorescence (IF) stainings

For standard IF stainings, cells were cultured on coverslips coated with 5 µg ml-1 fibronectin. Cells were fixed by 10 minute incubation with 4% paraformaldehyde in PBS++ (PBS with 1 mM CaCl_2_ and 0,5 mM MgCl_2_). Fixed cells were permeabilized for 5 minutes with 0,5% Triton X100 in PBS and blocked for 30 minutes in 2% bovine serum albumin (BSA) in PBS. Primary and secondary antibodies were diluted in 0,5% BSA in PBS and incubates for 45 minutes. After each step the fixed cells were washed three times with 0,5% BSA in PBS. Coverslips were mounted in Mowiol4-88/DABCO solution.

### Luciferase assays

HEK293 cells were seeded sparsely (75.000 cells per well) in 24-wells plates coated with gelatin. Cells were transfected with the pGL3-DLC1-promotor luciferase reporter plasmids using PEI (Polysciences). pRL-TK Renilla reporter plasmid was co-transfected (1/50) as a control for transfection efficiency. Two days after transfection Firefly and Renilla Luciferase activities were analyzed using the Dual-Luciferase Reporter Assay System (Promega) and the GloMax-Multi detection system (Promega) according to manufacturer’s protocol.

### Quantitative PCR (qPCR)

Total RNA was isolated from HUVECs using TRI Reagent (Sigma). cDNA synthesis was performed using iScript (Bio-Rad). Quantitative polymerase chain reaction was performed using SensiFAST SYBR Green No-ROX (Bioline) on a LightCycler 480 system (Roche). To calculate the relative gene expression, the ΔΔCt method was used. DLC1 expression was corrected for RPLP0 reference expression, and DLC1 expression levels were normalized to its levels on plastic. Primer sequences were as follows (5’-3’): DLC1 forward ATGATCGCCGAGTGCAAGAA and reverse CTGCTCCGAAGTGGAGTAGC. RPLP0 forward TCGACAATGGCAGCATCTAC and reverse ATCCGTCTCCACAGACAAGG.

### Scratch assays

For scratch assays, HUVECs were plated on 12-well or 24-well plates coated with 5 µg ml-1 fibronectin. Two perpendicular scratches per well were made using a sterile 200 µl pipette tip. Next, cells were washed with PBS^++^ supplied with EGM-2 medium, and were mounted on an inverted NIKON Eclipse TI microscope equipped with Okolab cage incubator and humidified CO_2_ gas chamber set to 37°C and 5% CO_2_. Cells were live imaged (16-20 hours; 10 min time interval) using phase-contrast imaging using a 10x CFI Achromat DL dry objective (N.A. 0.25) and an Andor Zyla 4.2 plus sCMOS camera. Images were enhanced for display with an unsharp mask filter and adjusted for brightness/contrast in ImageJ. Scratch wounding surface was quantified by measuring the wound area using the freehand tool in ImageJ. The ImageJ manual tracking plugin was used for single cell tracking, and the Chemotaxis tool was used to quantify directionality and velocity. For IF stainings of scratch assays, cells were plated on coverslips coated with 5 µg ml-1 fibronectin and fixed after 6 hours with 4% PFA. Golgi orientation was assessed by measuring the center of mass of the DAPI and GM130 signal and calculating the angle between these points in relation to the direction of migration. Focal adhesion orientation was analyzed using the focal adhesion server using a minimal adhesion size of 4 pixels and a maximal adhesion size of 115 pixels (Berginski and Gomez, 2013).

### Sprouting angiogenesis assays

For sprouting angiogenesis assays HUVECs were resuspended in EGM-2 medium containing 0.1% methylcellulose (4000 cP, Sigma). For spheroid formation 750 cells per 100 □l methylcellulose medium were seeded in wells of a U-bottom 96-wells plate and incubated overnight. Spheroids were collected and resuspended in 1,7 mg/ml collagen Type I rat tail mixture (IBIDI), plated in glass-bottom 96 well plates and placed at 37°C. After polymerization of the collagen gel, spheroids were stimulated with 50 ng ml-1 VEGF to induce sprouting overnight as described previously (Korff and Augustin, 1999). Pictures were taken using the EVOS M7000 imaging system and a 10x objective. Images were enhanced for display with an unsharp mask filter and adjusted for brightness/contrast in ImageJ. Sprouting number and length was analyzed using the ImageJ plugin NeuronJ.

### Fluorescence microscopy

For live cell fluorescence microscopy cells were plated on Lab-Tek chambered 1.0 borosilicated coverglass slides coated with 5 µg ml-1 fibronectin and cultured in EGM2 culture medium. For TIRF microscopy we used an inverted NIKON Eclipse TI equipped with a 60x 1.49 NA Apo TIRF (oil) objective, perfect focus system, Orange Diode Solid State Laser 594nm 30mW (Excelsior, Spectra-physics), dual band 488/594 nm TIRF filter cube (Chroma TRF59905 ET), and an Andor Zyla 4.2 plus sCMOS camera (without binning). An Okolab cage incubator and humidified CO_2_ gas chamber set to 37°C and 5% CO2 were used during the imaging process. Images was performed every 30 second interval for 3-5 hours. To analyze focal adhesion dynamics the raw data was uploaded to the focal adhesion server using a minimal adhesion size of 4 pixels and phase length of 5 minutes (Berginski and Gomez, 2013). For widefield microscopy of immunostained HUVECs (Fig. 3f, 3g, 4c, 5d) the NIKON Eclipse TI was equipped with a lumencor SOLA SE II light source and standard DAPI, GFP, mCherry or Cy5 filter cubes (NIKON). IF stained samples in Fig. 2b and 2e were imaged using an inverted Zeiss widefield microscopes Observer.Z1 equipped with a 63x 1.40 Plan Apochromat oil objective and a Hamamatsu Orca-R2 digital camera. Images were enhanced for display with an unsharp mask filter and adjusted for brightness/contrast in ImageJ.

### Traction Force Microscopy

For traction force microscopy HUVECS were plated on collagen coated 1.2 kPa (Young’s modulus) polyacrylamide substrates containing 2 □m reference bottom beads and 0.2 □m sulfated top beads (FluoSpheres, Molecular Probes). HUVECS were cultured to confluency on the gels for 48 hours and subsequently visualized using an inverted Zeiss Axiovert 200 widefield microscope equipped with a 40x 0.75 NA Zeiss air objective, Cooke Sensicam CCD camera and IBIDI climate-control system. To determine traction forces the top and reference beads were imaged using fluorescence microscopy and DIC to visualize the HUVECs. Finally, the HUVECs were trypsinized from the substrate to acquire images of the unloaded fiducial bead patterns. Computation of traction forces was performed as described previously using known material properties (stiffness=1.2 kPa, Poisson’s ration=0.48) and constrained twodimensional fast Fourier transformation method (Valent et al., 2016). From the monolayer traction fields, the root mean squared value of traction in Pascal was calculated as scalar measure for monolayer traction.

### Western blot analysis

Cells were lysed with reduced sample buffer containing 4% β-Mercaptoethanol. Samples were boiled at 95° for 5-10 minutes to denature the proteins. 10% SDS-page gels were used in 10X SDS-Page running buffer (250 mM Tris, 1,92 M glycine, 1% SDS, pH 8,3), and transferred to ethanol activated PVDF membrane using wet transfer in blot buffer (25 mM Tris, 192 mM glycine, 20% ethanol (v/v)). Blots were blocked with 5% milk powder in Tris-buffered saline (TBS) for 30 minutes. Blots were incubated overnight at 4°C with the primary antibodies in 5% ELK in TBS with Tween20 (TBST). The secondary antibodies linked to horseradish peroxidase (HRP) were incubated for 45 minutes at room temperature. As final step before imaging blots are washed with TBS. HRP signals were visualized by enhanced chemiluminescence (ECL) detection (SuperSignal West Pico PLUS, Thermo Fisher, Cat # 34580) and visualized on the ImageQuant LAS 4000 (GE Healthcare). Intensities of bands were quantified using Gel Analyzer plugin in ImageJ.

## Acknowledgements

This study was financially supported by the Netherlands Organization of Scientific Research (NWO-VIDI grant 016.156.327).

---

**Supplemental Figure 1.**
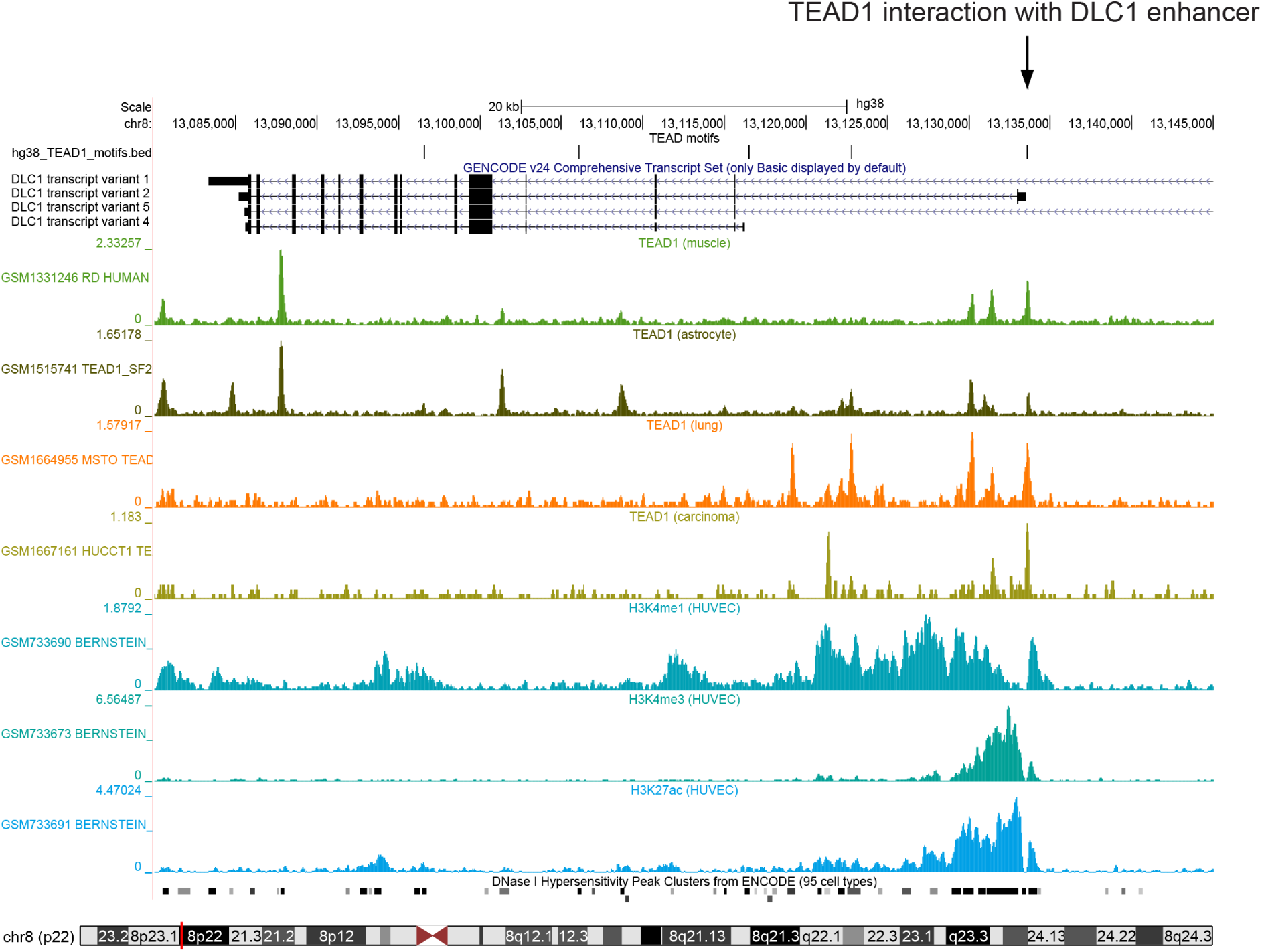
Full detailed schematics of UCSC genome browser results at position chr8:13,074,715-13,142,890 of the human genome (GRCh38/hg38) showing the genomic location of DLC1 transcript variants 1 (NP_872584.2), 2 (NP_006085.2), 5 (NP_001303597.1) and 4 (NP_001157743.1) and the presence of a TEAD motif at the transcriptional start site (TSS) of DLC1 transcript variant 2. Plotted are the results from publicly available GEO data TEAD1 ChIP-Seq data from various cell types and corresponding histone modification profiles in HUVECs in ENCODE. The data show a binding peak of TEAD1 at the TSS of DLC1 transcript variant 2. Histone modification profiles indicate that there is an open conformation of chromatin and an active promoter region around the TEAD binding motif, defined as the bimodal presence of both histone H3 trimethylation at lysine 4 (H3K4me3) and histone H3 acetylation at lysine 27 (H3K27ac), combined with increased DNase hypersensitivity.

**Supplemental Figure 2.**
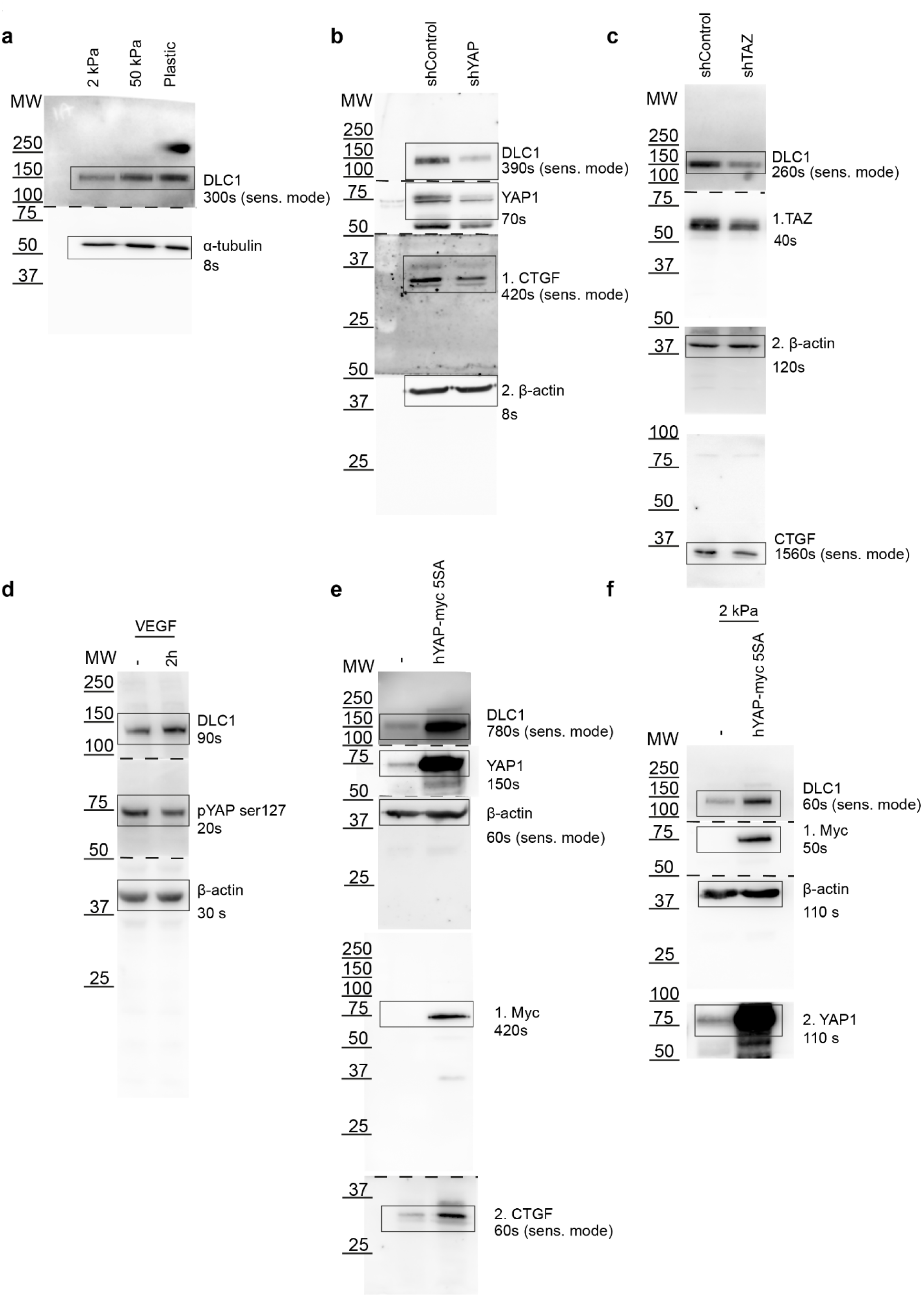
Full scans of Western experiments in Figure 1. Molecular weights of the marker, exposure times, sensitive scanning mode and following order of antibody probing are indicated.

**Supplemental Figure 3.**
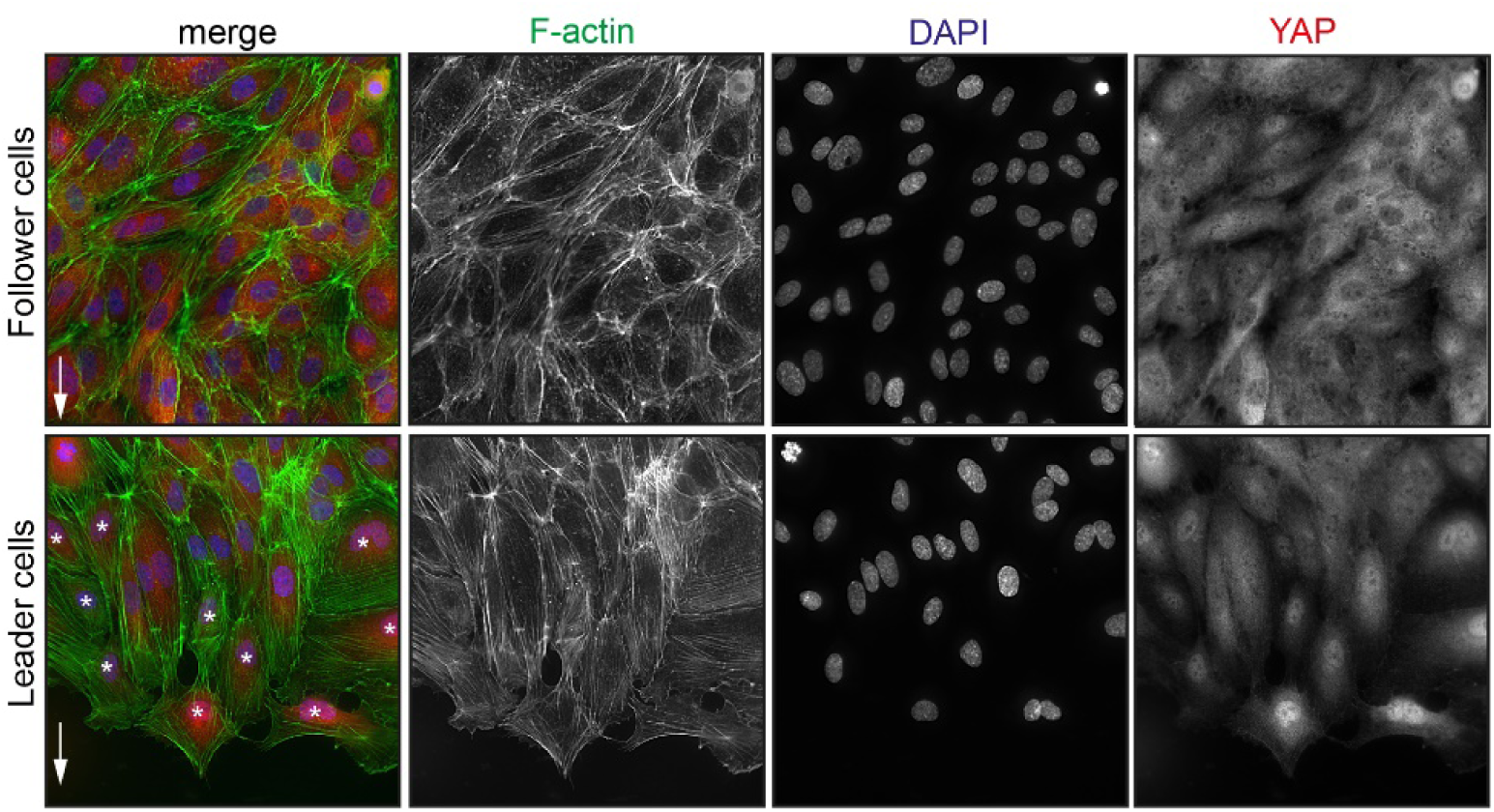
Widefield IF images of HUVECs fixed 6 hours after initiation of scratch wound assays stained for DAPI (blue), F-actin (green) and YAP (red). Pictures taken of the follower cells in the center of the monolayer and of the leader cells at the scratch wound edge. Asterisks highlight cells with nuclear enrichment of YAP compared to thecytoplasm.

**Supplemental Figure 4.**
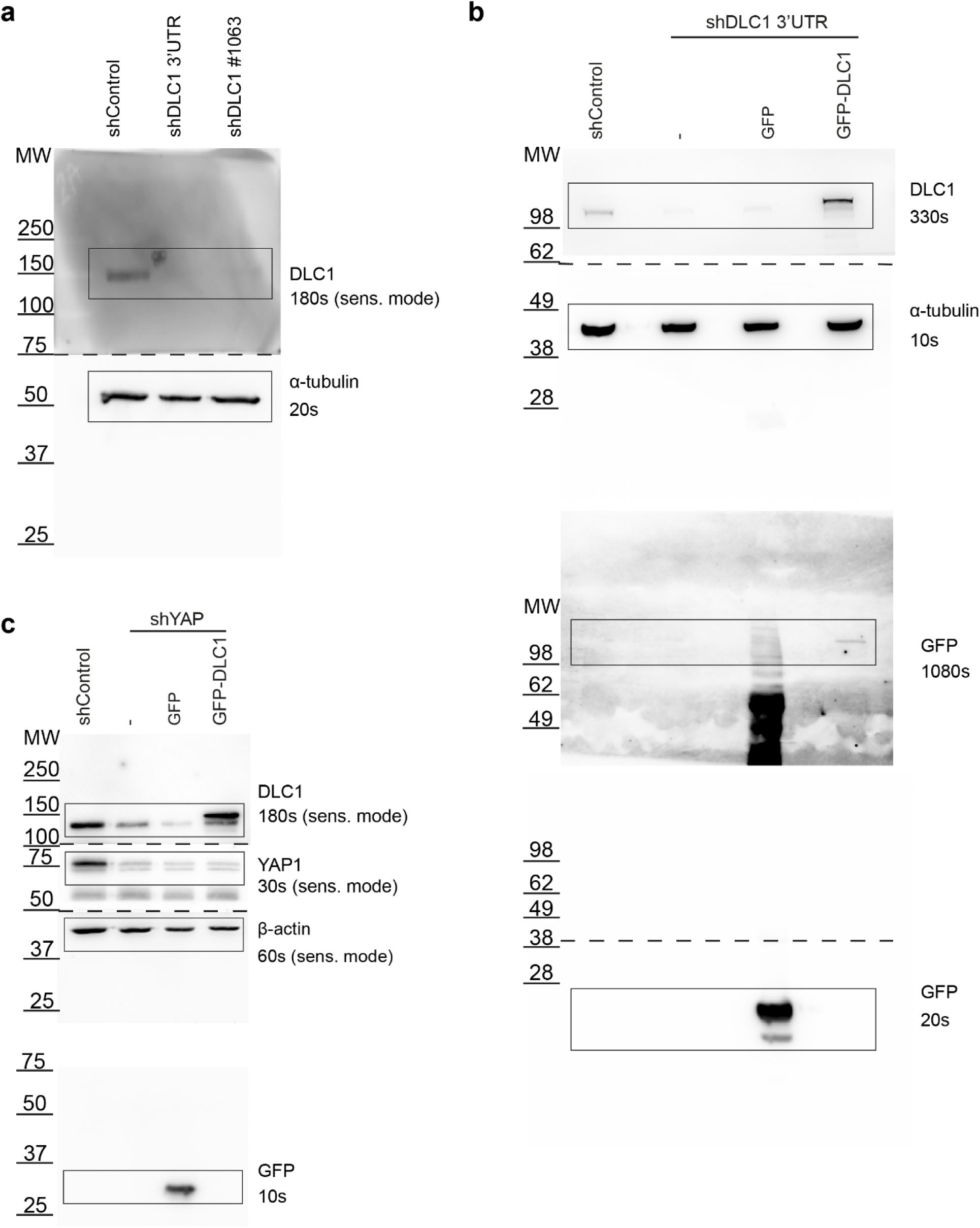
Full scans of Western experiments in Figure 3-5. Molecular weights of the marker, exposure times, sensitive scanning mode and following order of antibody probing are indicated.

**Supplemental Movie 1. DLC1 controls endothelial focal adhesion dynamics.** Time lapse recording of HUVECs transduced with shControl or shDLC1 and expressing paxillin-mCherry. Images were acquired by time-lapse TIRF microscopy (NIKON Eclipse Ti) using a 60x/1.49 NA oil objective. Frames were taken every 30 sec for ∼ 2,5 hours.

**Supplemental Movie 2. DLC1 is needed for endothelial directional migration.** Time lapse recording of HUVECs transduced with shControl, shDLC1 3’UTR or shDLC1 #1063 during scratch wound migration. Images were acquired by time-lapse phase-contrast microscopy (NIKON Eclipse Ti) using a 10x dry objective. Frames were taken every 10 min for ∼ 16 hours.

